# Predicting assembly/disassembly orders of protein complexes using coarse-grained simulations

**DOI:** 10.1101/2024.01.24.576999

**Authors:** Yunxiao Lu, Xin Liu, Zhiyong Zhang

**Author notes:** Corresponding Author(s): Zhiyong Zhang.

## Abstract

Assembly of a protein complex is very important to its biological function, which can be investigated by determining assembly/disassembly order of its protein subunits. Although static structures of many protein complexes are available in the protein data bank, their assembly/disassembly orders of subunits are largely unknown. In addition to experimental techniques for studying subcomplexes in the assembly/disassembly of a protein complex, computational methods can be used to predict the assembly/disassembly order. Since sampling is a nontrivial issue in simulating the assembly/disassembly process, coarse-grained simulations are more efficient than atomic simulations are. In this work, we developed computational protocols for predicting assembly/disassembly orders of protein complexes using coarse-grained simulations. The protocols were illustrated using two protein complexes, and the predicted assembly/disassembly orders are consistent with available experimental data.

## Introduction

Protein complexes carry out the majority of catalytic, structural, and regulatory functions^[1]^. Currently, cryo-electron microscopy (cryo-EM)^[2]^ and X-ray crystallography^[3]^ are the primary experimental techniques for obtaining high-resolution atomic structures of protein complexes. Additionally, computer techniques incorporating artificial intelligence enable predictive modeling of protein complexes^[4]^. With the continuous advancement of experimental and computational methods, an increasing number of structures of protein complexes have been determined, providing opportunities to investigate their functions at the molecular level.

However, in addition to static structures, it is essential to study assembly of protein complexes^[5]^. Many protein complexes have evolved to be assembled in a specific order^[6]^. The stability and function of a protein complex rely on a multistep process of subunit assembly, which may provide valuable insights into the function and evolution of the protein complex. Numerous studies have demonstrated that improper protein assembly may result in negative biological effects^[7]^. However, due to the high complexity of the assembly pathway, the current understanding of the assembly mechanism is limited. According to Tompa and Rose’s study^[8]^, the assembly is hierarchical, and a protein complex must follow an ordered assembly pathway to achieve its biological function^[9]^. Therefore, studying the assembly of protein complexes contributes to the reconstruction of protein complexes in vitro, the design of artificial protein complexes, and the development of new drugs related to assembly^[10]^.

The assembly of a protein complex can be studied by determining assembly/disassembly order of its subunits. Stabilized intermediates of two or more subunits are detected using experimental techniques such as co-immunoprecipitation^[11]^, mass spectrometry (MS)^[12]^, and time-resolved cryo-electron microscopy^[13]^, or by creating deletion mutants to study the effect of deletion on assembly^[14]^. MS is a powerful technique for identifying subcomplexes during the assembly/disassembly of a protein complex^[15]^. A typical MS measurement generally starts with ionization of the protein complex^[16]^. The complex is excessively charged, and signals from a set of subcomplexes are subsequently detected via MS. Subunits that bind each other weakly tend to dissociate from the complex^[17]^.

Computational modeling has the potential to predict the assembly/disassembly order of protein complexes. In early studies^[18, 19]^, the order of subcomplex formation was predicted using a simple model based on interface size, which was given by the number of contacted residues^[20]^. With seven out of nine heteromeric complexes, they found agreement between interface sizes and assembly/disassembly orders. Path-LZerD^[21]^ uses knowledge-based potentials to evaluate binding preferences between subunits and can predict assembly paths even when the complex structure is unknown. The method involves constructing docked structures of subunits from single subunits using the Multi-LZerD method^[22]^, and iterative optimizations are performed via a genetic algorithm^[23]^. The predictions of Path-LZerD are consistent with the experimental data for three complexes with up to four subunits, but for larger complexes with five or more subunits, the number of correct predictions decreased. The hybrid Monte Carlo/molecular dynamics simulation (hMC/MD) method^[24]^ uses a likelihood-based selection scheme to predict the disassembly order of a protein complex. The success rate of the method was greater than 0.9 for four tetrameric protein complexes.

When modeling large protein complexes, all-atom simulations are computationally expensive. Therefore, other methods, such as coarse-grained (CG) simulations, which are computationally efficient, have been proposed^[25]^. Although a CG model does not describe protein interactions as accurately as an all-atom model, the former can still capture kinetic features of protein complexes^[26]^. Furthermore, CG simulations can depict a biological process over a long timescale and have been employed to a variety of protein complexes^[25-27]^.

This work developed protocols for predicting assembly/disassembly order of protein complexes using CG simulations. In one protocol, the disassembly process of a protein complex is simulated via CG simulations at different temperatures. This idea comes from MS experiments because standard electrospraying relies on the use of some factors like high temperatures. The temperature can be used to simulate the degree of ionization in MS and induce the dissociation of protein complexes. The disassembly temperature of each subunit is estimated through multi-temperature CG simulations, after which the disassembly order of the complex is predicted. In the other protocol, the assembly order of a protein complex is obtained by running iterative CG simulations at the room temperature, in order to sequentially determine the most stable dimer, trimer, …, until the whole complex is assembled. Two protein complexes, the phosducin–Gtβγ complex and the Arp2/3 complex, were used to validate the protocols.

## Materials and methods

### The protocol for predicting the disassembly order

To predict the disassembly order of multiple subunits in a protein complex, a protocol based on multi-temperature CG simulations is used (**Fig. 1**). A high temperature breaks interactions between protein subunits, which may lead to subunit disassembly. This process can be more easily observed by CG simulations than by all-atom simulations.

**Figure 1.**
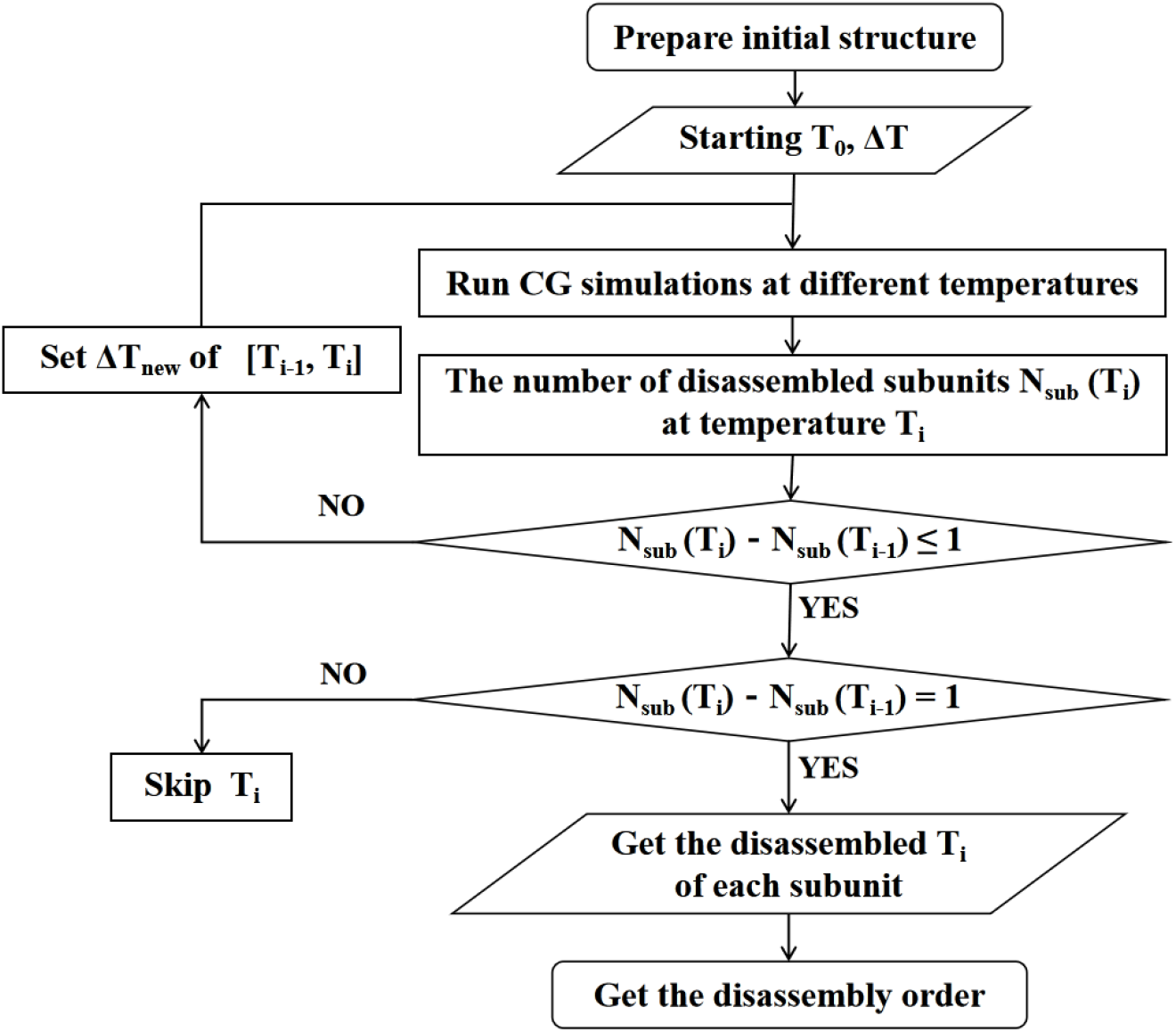
Flowchart of multi-temperature CG simulations for predicting the disassembly order of a protein complex.

The protocol consists of the following steps:

1. The initial structure of the protein complex is prepared, such as building missing regions.
2. CG simulations are run at different temperatures. The temperatures are 300 K, 300 K + ΔT, 300 K + 2ΔT, …, until all the subunits are disassembled at a certain temperature.
3. The CG trajectories are analyzed from low to high temperatures, and N_sub_ and ΔN_sub_ at each temperature are obtained. N_sub_(T_i_) is the number of disassembled subunits at temperature T_i_ that can be obtained by analyzing the qscore in the CG simulation (details can be found in the next section). ΔN_sub_(T_i_) is the difference between N_sub_ at temperature T_i_ and temperature T_i-1_.

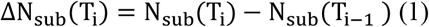
4. If ΔN_sub_(T_i_)=0, the temperature T_i_ is skipped. If ΔN_sub_(T_i_)=1, there is a disassembled subunit between T_i-1_ and T_i_, and we set the disassembled temperature of this subunit as T_i_. If ΔN_sub_(T_i_)>1, there are two or more disassembled subunits between T_i-1_ and T_i_. In this case, a ΔT_new_ needs to be set in the temperature range [T_i-1_, T_i_]. Similar to step 2, a series of CG simulations are run at T_i-1_+ΔT_new_, T_i-1_+2ΔT_new_, …, until T_i_-ΔT_new_. These CG trajectories are analyzed as described in step 3.
5. The procedure is repeated until the disassembled temperature of each subunit was determined.

### Determining the disassembly of a subunit by qscore

Native contacts are formed by pairs of amino acid residues that are physically close to each other in the native structure. The cutoff distance of the C_α_ atoms between two residues in native contact was set to 6.5 Å by default. The intersubunit native contacts in the complex were all counted.

qscore is the fraction of native contacts in a conformation. In the native structure of a protein complex, if two subunits have native contacts, the corresponding intersubunit qscore is 1.0. When the intersubunit qscore between two subunits is 0, there is no native contact between them. During a CG simulation, the qscores may change, which can be used to determine whether each subunit disassembles. If a subunit loses contact with other subunits (qscores become 0) in the CG simulation, the subunit is disassembled.

### The protocol for predicting the assembly order

To predict the assembly order of a protein complex, a protocol is developed by running iterative CG simulations of different subcomplexes, from dimers, trimers, …, until the intact complex is formed (**Fig. 2**).

**Figure 2.**
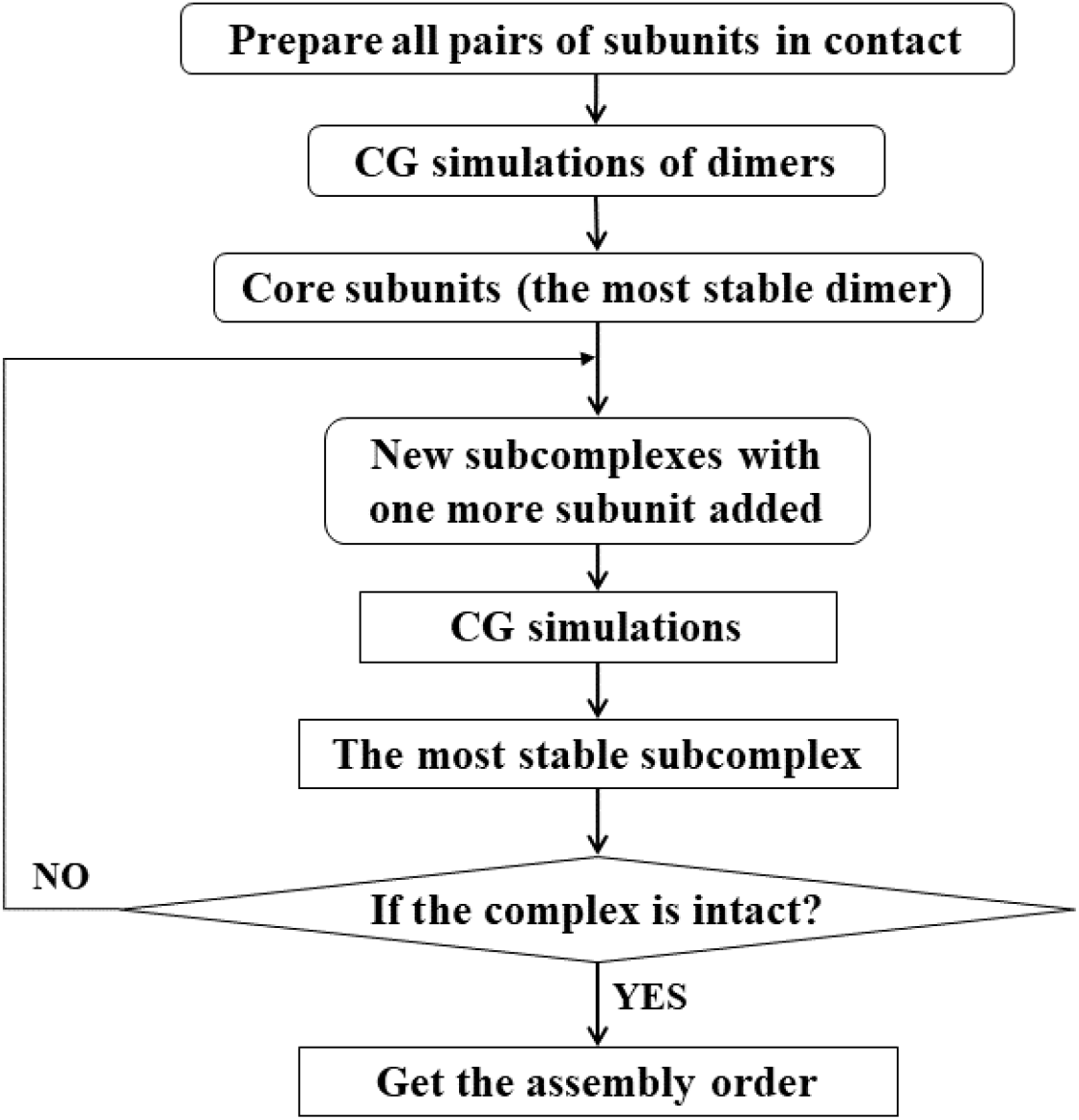
Flowchart of iterative CG simulations for predicting the assembly order of a protein complex.

The protocol consists of the following steps:

1. From the intact structure of the protein complex, all pairs of subunits in contact are prepared.
2. CG simulations at the room temperature (300 K) are run for all the dimers individually.
3. The most stable dimer is found. For example, one can calculate the average root mean square deviation (RMSD) of each dimer during the CG simulation, and pick the one with the smallest RMSD. The two subunits in this dimer are defined as core subunits.
4. Starting from the most stable dimer, another subunit in contact with the dimer is added to form a trimer. All the trimers are prepared.
5. CG simulations are run for all the trimers individually.
6. The most stable trimer is found.
7. Steps 4 to 6 are repeated until the intact protein complex is formed. The assembly order is finally determined.

It should be noted that this protocol may require a lot of CG simulations for all the possible subcomplexes. However, the core subunits can sometimes be defined by other means, and the steps 1 to 3 could be skipped.

### The CG Model

An atomic interaction-based CG model called AICG2+ was used for proteins^[28]^. The energy function of the AICG2+ model is as follows:

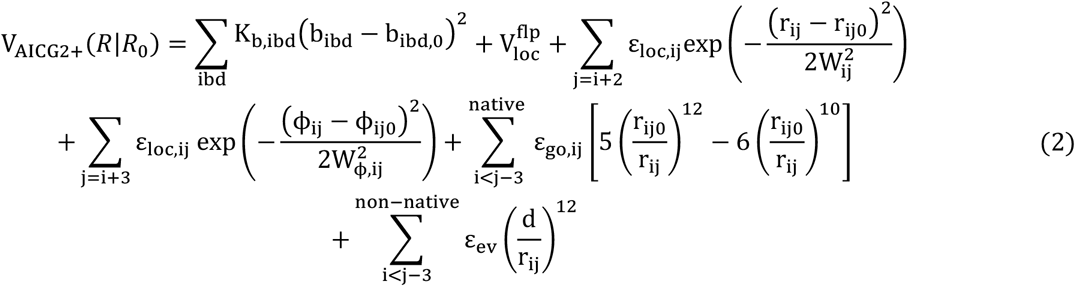

*R* represents the 3n_aa_ Cartesian coordinates of the simulated protein, and *R*_*0*_ represents those of the native structure. b_ibd_ is the ibd-th virtual bond length. r_ij_ is the distance between the *i*-th and *j*-th amino acids. 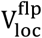 is a generic flexible local potential for virtual bond angles and dihedral angles constructed by analyzing loop structures in a protein structure database. The third term describes specific local interactions, and ε_loc,ij_ and W_ij_ in the Gaussian function represent the strength and width of the local interaction between residues i and j, respectively. Φ_ij_ in the fourth term is the dihedral angle formed by the four consecutive residues i to i+3, and W_Φ,ij_ is the width of this local interaction. The fifth term is the structure-based native contact potential, and the last term represents the sum of interactions between non-native contact residue pairs.

### CG Simulations

All CG simulations were run using the CafeMol 3.2 package, which is a general-purpose CG simulation software for simulating proteins and their complexes^[29]^. We used the AICG2+ model, excluded volume and electrostatic interactions for protein complexes. The electrostatic interactions were calculated using the Debye–Hückel equation with a dielectric constant of 78 F/m and an ion concentration of 0.15 M.

For each system, an energy minimization was initially performed at 300 K, running 1000 steps with the steepest descent method, followed by 2000 steps with the conjugate gradient algorithm. Subsequently, a production simulation was run using Langevin dynamics at a certain temperature. The time step was 0.4 Cafe-time (1 Cafe-time ∼ 49 fs). In total, 10^8^ steps were run for each simulation, and a snapshot was saved every 5000 steps. The neighbor list was updated every 100 steps.

The parameter go_unit determines the weight of the Gō potential in the energy function, and its default value 1.0 may make the complex rigid. Usually, different go_unit from 0.5 to 1.0 are tested by running CG simulations at 300 K. Finally a relatively small go_unit is chosen, with which the complex is not rigid in the CG simulation while essentially keeping stable.

### Protein complexes for testing the methods

We selected two asymmetric heteromeric protein complexes from the 3D Complex server^[20]^, the phosducin– Gtβγ complex (**Fig. 3a)** and the Arp2/3 complex (**Fig. 3b**), for testing the protocols.

**Figure 3.**
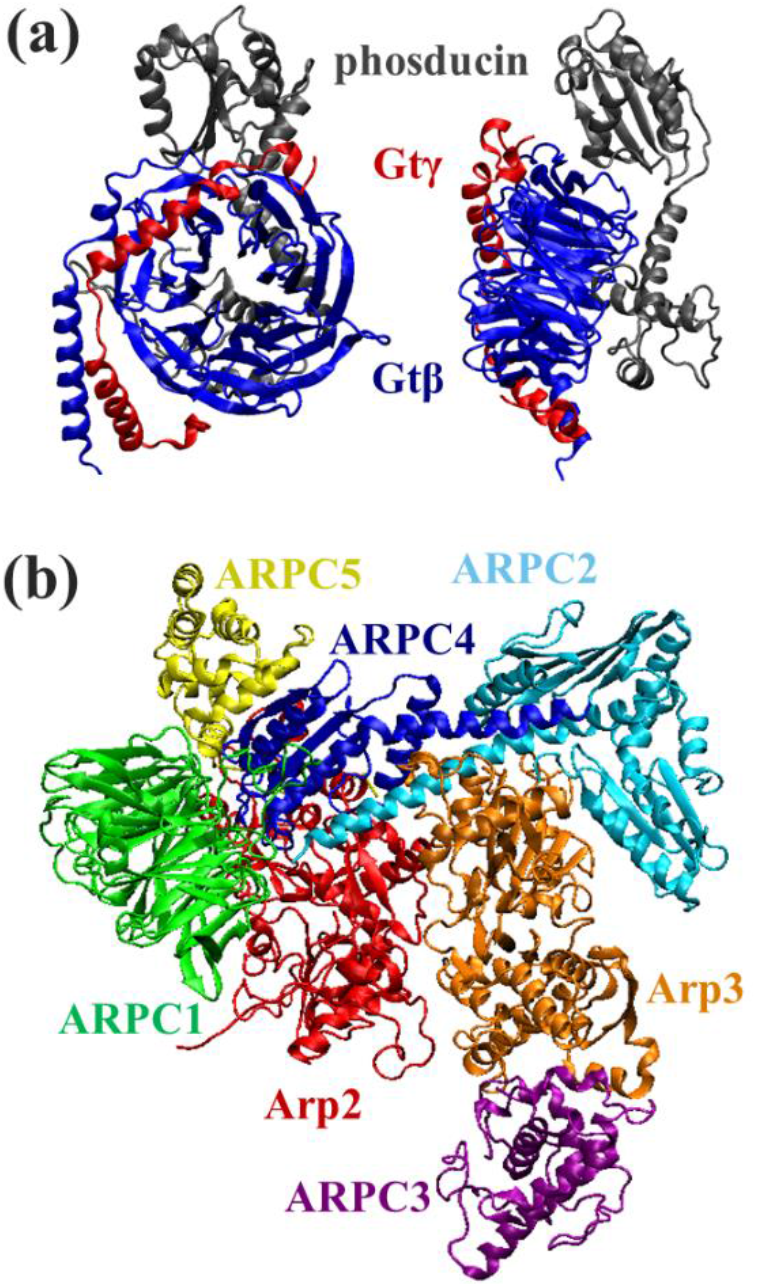
Structures of the two protein complexes used to test the protocols. (a) The phosducin–Gtβγ complex. (b) The inactive state of the Arp2/3 complex. The ATP/ADP molecules were excluded in the CG simulations.

### The phosducin–Gtβγ complex

Phosducin binds tightly to Gtβγ of the heterotrimeric G protein transducin, preventing Gtβγ reassociation with Gtα–GDP and thereby inhibiting the G-protein cycle^[30]^. The crystal structure of the phosducin–Gtβγ complex (PDB 1A0R) from *bovine retina* was solved with a resolution of 2.80 Å^[31]^. The complex consists of three subunits, phosducin and the Gtβγ dimer (**Fig. 3a**). The β subunit forms a seven-bladed β propeller around which the γ subunit is wrapped, with the N-terminal helices of the β and γ subunits forming a coiled coil. Phosducin is composed of two domains: a helical domain at the N-terminus and a mixed αβ domain at the C-terminus. The former covers the top of the β propeller, while the latter covers one of its sides^[32]^. From experimental data, the Gtβγ dimer can be formed in the absence of phosducin, therefore, the order of assembly is known^[33]^.

### The Arp2/3 complex

This complex is an important regulator of the cytoskeleton and a factor that enhances the mobility of cancer cells, showing a strong correlation with cancer development^[34]^. The crystal structure of the inactive Arp2/3 complex (PDB 1K8K) from *bos taurus* was solved with a resolution of 2.0 Å^[35]^, and the non-terminal missing residues were built by homology modeling^[36]^. The complex consists of seven subunits, including two actin-associated proteins (Arp2 and Arp3) and five other proteins (named ARPC1-ARPC5 by descending order of molecular mass)^[35]^. Both Arp2 and Arp3 are structurally similar to actin and may form a dimer with the ability to bind actin filaments to form branches^[37]^. ARPC2 and ARPC4 hold the dimer together by anti-parallel binding of the C-terminal long helix(**Fig. 3b**). The α/β domain of ARPC4 interacts with Arp2, ARPC1, and ARPC5, while the two similar α/β domains of ARPC2 interact only with Arp3^[38]^. ARPC3 is only in contact with Arp3^[39]^.

## Results and Discussion

### Disassembly order of the phosducin–Gtβγ complex

First, we obtained a matrix of native contacts of the phosducin–Gtβγ complex (Table 1) using CafeMol^[29]^. Gtβ and Gtγ have the largest number of intersubunit contacts, which is consistent with the experimental data showing that the two subunits serve as the dimer of the transducin

**Table 1.**
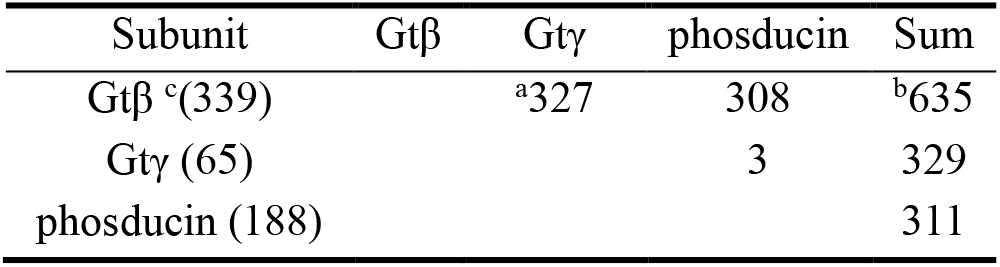
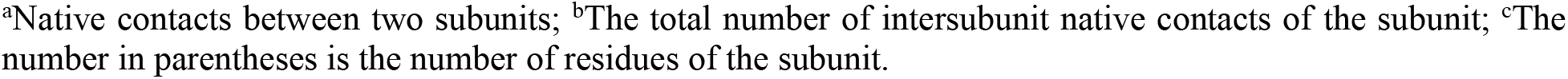
Intersubunit native contacts in the atomic structure of the phosducin–Gtβγ complex.

| Subunit | Gtβ | Gtγ | phosducin | Sum |
| --- | --- | --- | --- | --- |
| Gtβ <sup>c</sup>(339) |  | <sup>a</sup>327 | 308 | <sup>b</sup>635 |
| Gtγ (65) |  |  | 3 | 329 |
| phosducin (188) |  |  |  | 311 |

<sup>a</sup>Native contacts between two subunits; <sup>b</sup>The total number of intersubunit native contacts of the subunit; <sup>c</sup>The number in parentheses is the number of residues of the subunit.

The go_unit was set to 0.65. This value would allow for a significant conformation change of the complex in a CG simulation at 300 K, while the complex structure can be still preserved. Then, we performed independent CG simulations under different temperatures with the scheme shown in Figure 1 to simulate the disassembly processes of the complex.

For each subunit i, a weighted average of inter-subunit qscores (denoted as <inter-q>_i_) was computed as the follow:

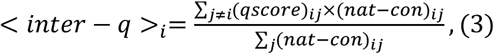

where (qscore)_ij_ is the qscore between subunits i and j, and (nat-con)_ij_ is the number of native contacts between the two subunits. The time evolution of <inter-q> during the CG simulation at each temperature was investigated.

In the CG simulation at 367 K (**Fig. 4**), the <inter-q> of phosducin (grey) drops to zero firstly, followed by Gtβ (blue) and Gtγ (red). The dissociation times of the three subunits are very close to each other. The structure of Gtβ is largely unfolded when phosducin disassembled, and then the β and γ subunits dissociates very shortly. Therefore, the predicted disassembly order of the phosducin-Gtβγ complex is phosducin, and the Gtβγ dimer.

**Figure 4.**
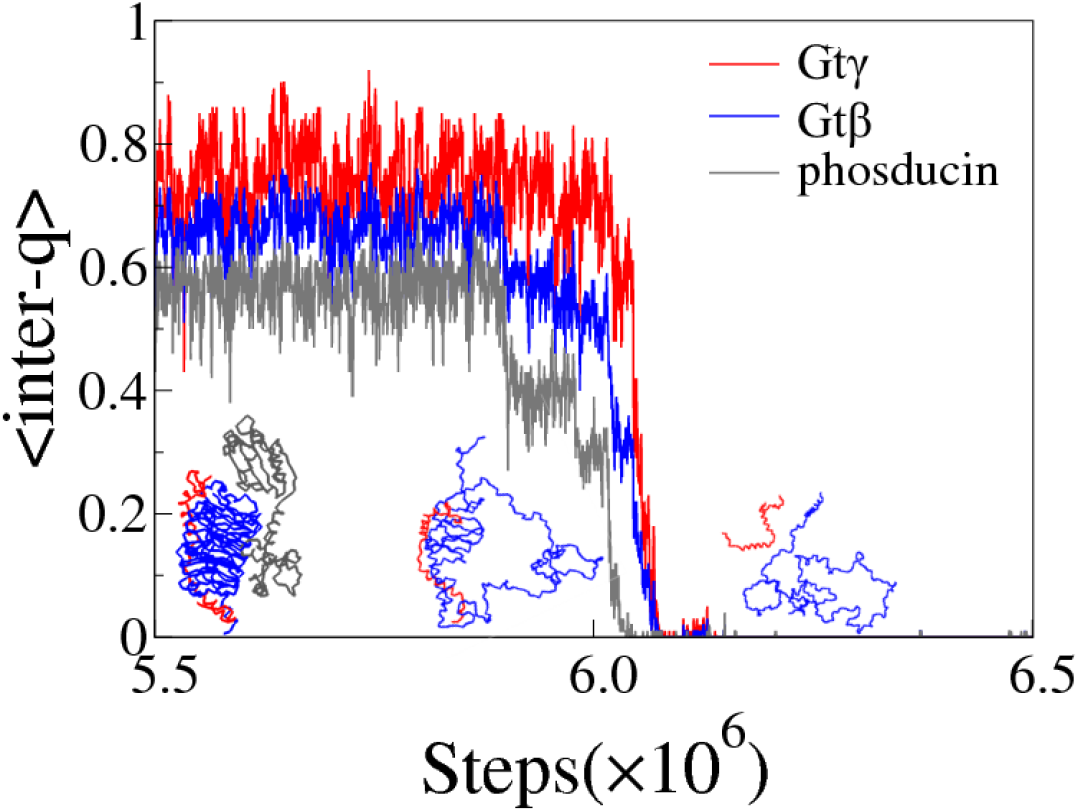
Time evolution of <inter-q> for every subunit in the phosducin–Gtβγ complex during the CG simulation at 367 K. Some snapshots are shown.

### Assembly order of the phosducin–Gtβγ complex

The assembly order of the phosducin–Gtβγ complex was investigated using iterative CG simulations of subcomplexes at 300 K (**Fig. 2**). First, CG simulations were performed to find the most dimer as the core subunits. Comparison of RMSD values of the three possible dimers indicates that the Gtβγ dimer is the most stable, with an average RMSD value of 2.0 Å (**Fig. 5**). In contrast, the phosducin-Gtγ dimer has a very large RMSD value (more than 400 Å) because there are few contacts between the two subunits (Table 1) and they dissociate even in the CG simulation at the room temperature. Therefore, the β and γ subunits assemble first to form the core dimer, followed by phosducin to form the trimer. Notably, our predicted assembly order is the inverse of the disassembly order predicted by CG simulations at 367 K (Fig. 4), which supports the notion that the assembly and the disassembly processes are generally reversible in protein complexes^[19]^.

**Figure 5.**
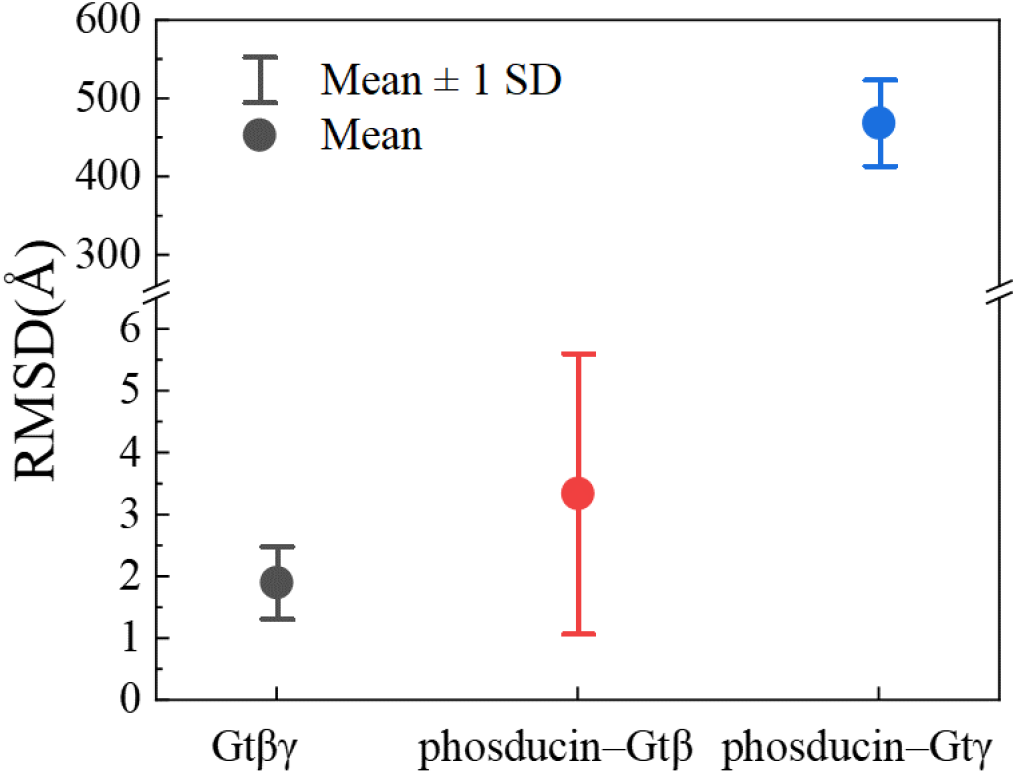
The mean and standard deviation of the RMSD value of dimers in the phosducin–Gtβγ complex. The RMSD value of each CG simulation was calculated using the last 100 conformations of the trajectory.

### Disassembly order of the Arp2/3 complex

Table 2 lists the matrix of native contacts of the Arp2/3 complex. ARPC2 and ARPC4 have the largest number of intersubunit contacts, which is consistent with the experimental data showing that ARPC2 and ARPC4 serve as the core subunits of the complex^[39]^.

**Table 2.** Inter-subunit native contacts in the atomic structure of the Arp2/3 complex.

| Subunit | ARPC4 | ARPC2 | Arp3 | Arp2 | ARPC1 | ARPC5 | ARPC3 | Sum |
| --- | --- | --- | --- | --- | --- | --- | --- | --- |
| ARPC4<br><sup>c</sup>(167) |  | <sup>a</sup>244 | 72 | 102 | 188 | 107 | 0 | <sup>b</sup>713 |
| ARPC2<br>(282) |  |  | 213 | 0 | 2 | 0 | 0 | 459 |
| Arp3<br>(414) |  |  |  | 79 | 0 | 0 | 88 | 452 |
| Arp2<br>(378) |  |  |  |  | 37 | 98 | 0 | 316 |
| ARPC1<br>(368) |  |  |  |  |  | 80 | 0 | 307 |
| ARPC5<br>(143) |  |  |  |  |  |  | 0 | 285 |
| ARPC3<br>(173) |  |  |  |  |  |  |  | 88 |

<sup>a</sup>Native contacts between two subunits; <sup>b</sup>The total number of inter-subunit native contacts of the subunit; <sup>c</sup>The number in the parenthesis is the number of residues of the subunit.

The go_unit was also set to 0.65, and CG simulations at different temperatures (**Fig. 1**) were performed to simulate the disassembly processes of the Arp2/3 complex.

At 315 K (**Fig. 6a**), the <inter-q> of Arp3 (orange), ARPC2 (cyan), and ARPC3 (purple) are essentially greater than 0.8. However, the <inter-q> of ARPC1 (green) drops to zero in the middle of the CG simulation, and then followed by ARPC5 (yellow). That is to say, ARPC1 and ARPC5 are disassembled sequentially at 315 K (**Fig. 7**). The <inter-q> of Arp2 (**Fig. 6a**, red) and ARPC4 (**Fig. 6a**, blue) are decreased since they have lost contacts with the disassembled ARPC1 and ARPC5. It has been found that Arp2 is unfolded after the disassembly of ARPC1 although the former is still in contact with the core (**Fig. 7**). In the CG simulation at 316 K (**Fig. 6b**), Arp2 is disassembled since its <inter-q> drops to zero. The <inter-q> of Arp3 and ARPC4 are decreased accordingly because they have lost contacts with the disassembled Arp2. The <inter-q> of ARPC2 and ARPC3 still keep at high values. The sub-complex ARPC2-ARPC4-Arp3-ARPC3 is well folded at this temperature (**Fig. 7**). At 350 K (**Fig. 6c**), ARPC3 is disassembled. It should be noted that, ARPC3 only has 88 native contacts with Arp3 (Table 2). However, the disassembly of ARPC3 requires a relatively high temperature, indicating that the binding of ARPC3 to Arp3 is strong. The <inter-q> of Arp3 is further decreased at 350 K because its contacts with ARPC3 have broken, and that of ARPC2 starts to decrease. It has been found that the sub-complex ARPC2-ARPC4-Arp3 is largely unfolded at this temperature (**Fig. 7**). Finally, in the CG simulation at 352 K, Arp3 is disassembled and then followed by the ARPC2-ARPC4 core (**Fig. 6d**). The core dimer (**Fig. 7**) is the last to disassemble.

**Figure 6.**
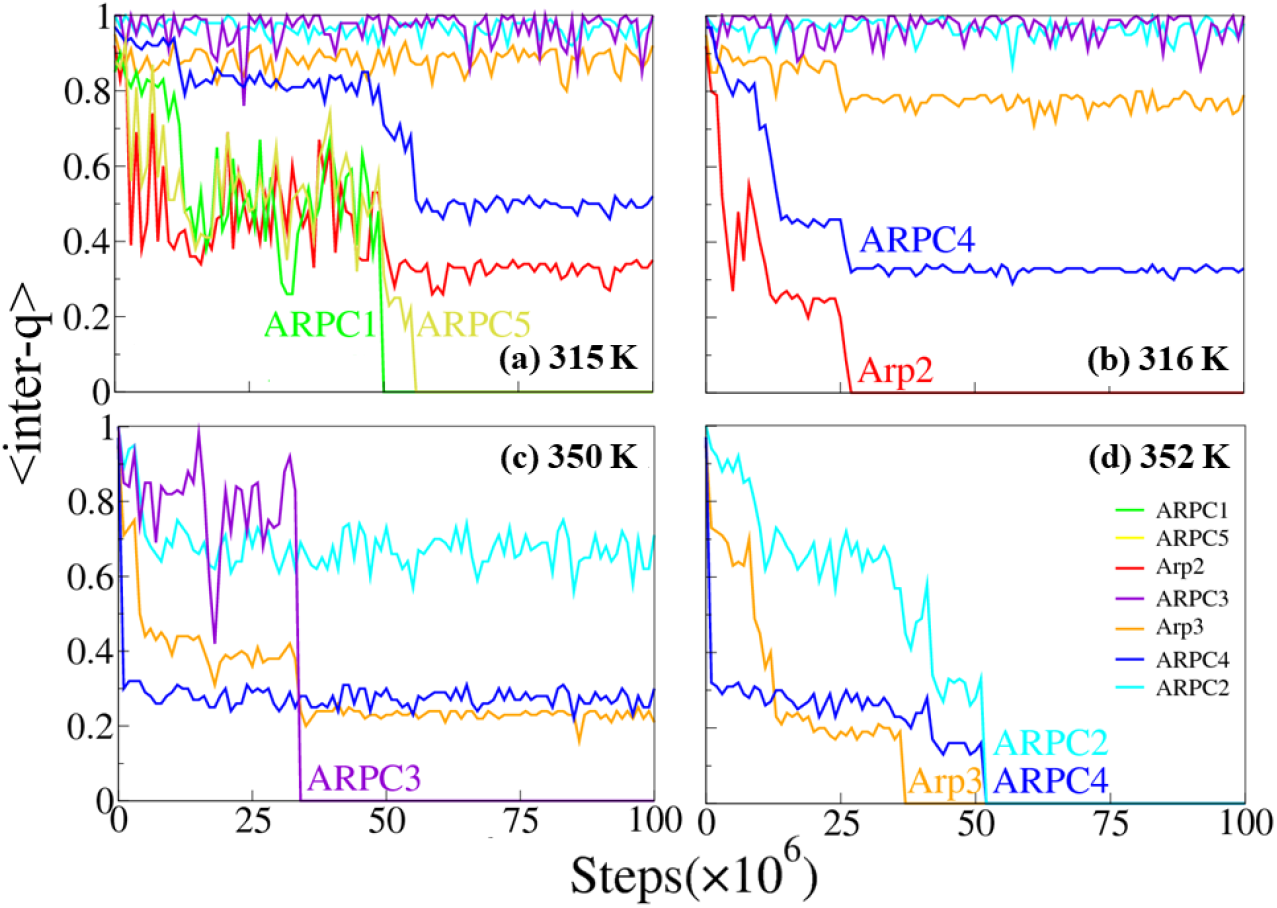
Time evolution of <inter-q> for every subunit in the Arp2/3 complex during the CG simulations at different temperatures. (a) 315 K, (b) 316 K, (c) 350 K, (d) 352 K.

**Figure 7.**
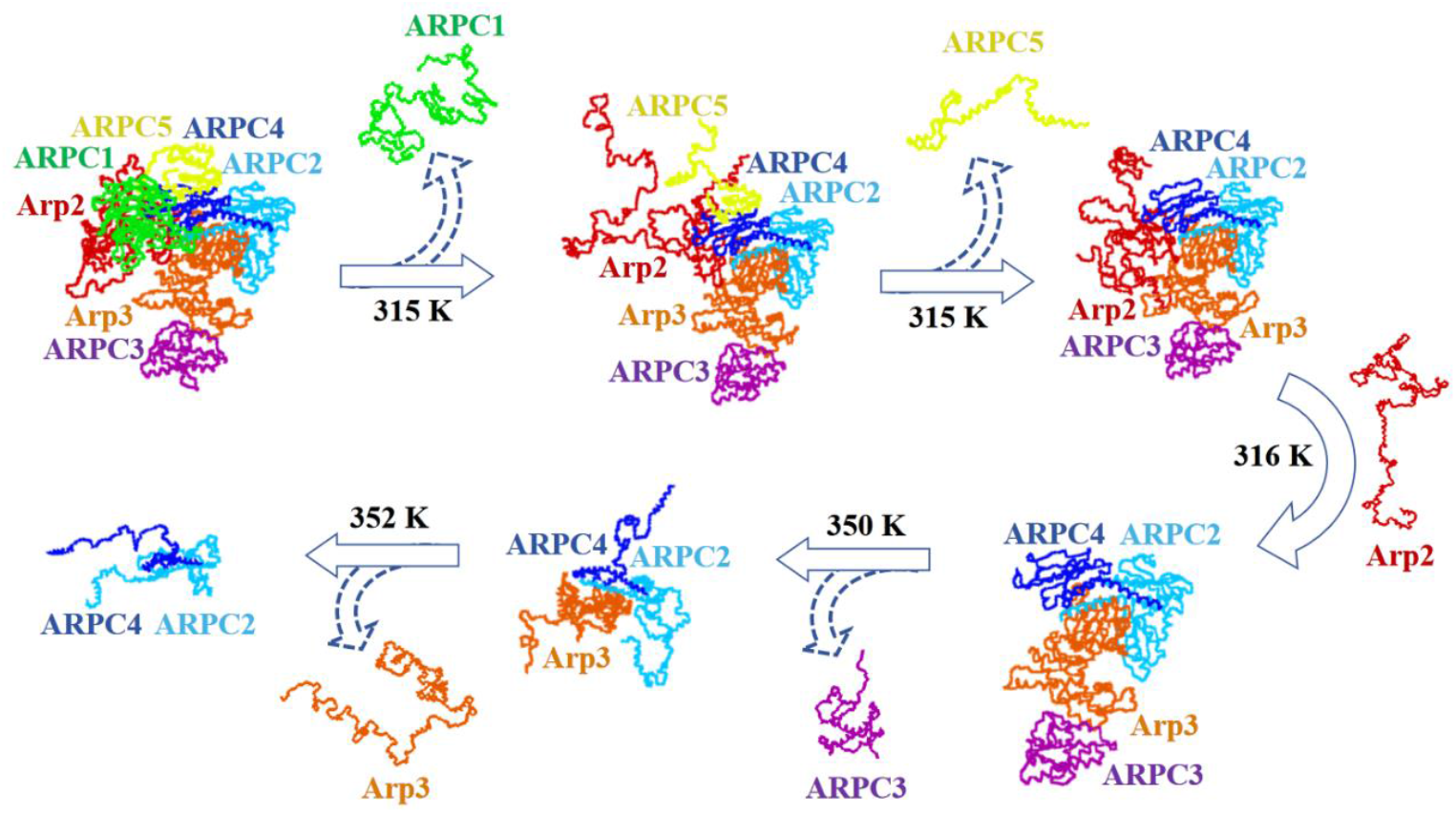
Snapshots during the disassembly process of the Arp2/3 complex at different temperatures.

According to the above analysis, the disassembly temperatures (order) of the peripheral subunits are 315 K (ARPC1 and ARPC5), 316 K (Arp2), 350 K (ARPC3), and 352 K (Arp3). If two or more subunits have the same disassembly temperature, we determine their disassembly order according to their <inter-q> plots (**Fig. 6**). The procedure was repeated three times independently, and the same disassembly order of the subunits was obtained.

### Assembly order of the Arp2/3 complex

The assembly order of the Arp2/3 complex was investigated using iterative CG simulations of subcomplexes (**Fig. 2**). Since ARPC2 and ARPC4 are known to be the core subunits, in the first iteration, only the CG simulation of the core dimer was conducted. The dimer is stable with an average RMSD of 2.1 Å.

In the second iteration, each of the peripheral subunits except ARPC3 was added to the core dimer, and the corresponding trimer was used to run a CG simulation. ARPC3 is not considered in this iteration because it has no direct contact with the core dimer. ARPC2-ARPC4-Arp3 is the most stable trimer, with the smallest average RMSD of 2.0 Å (**Fig. 8a**). Therefore, Arp3 may assemble first to the core dimer.

**Figure 8.**
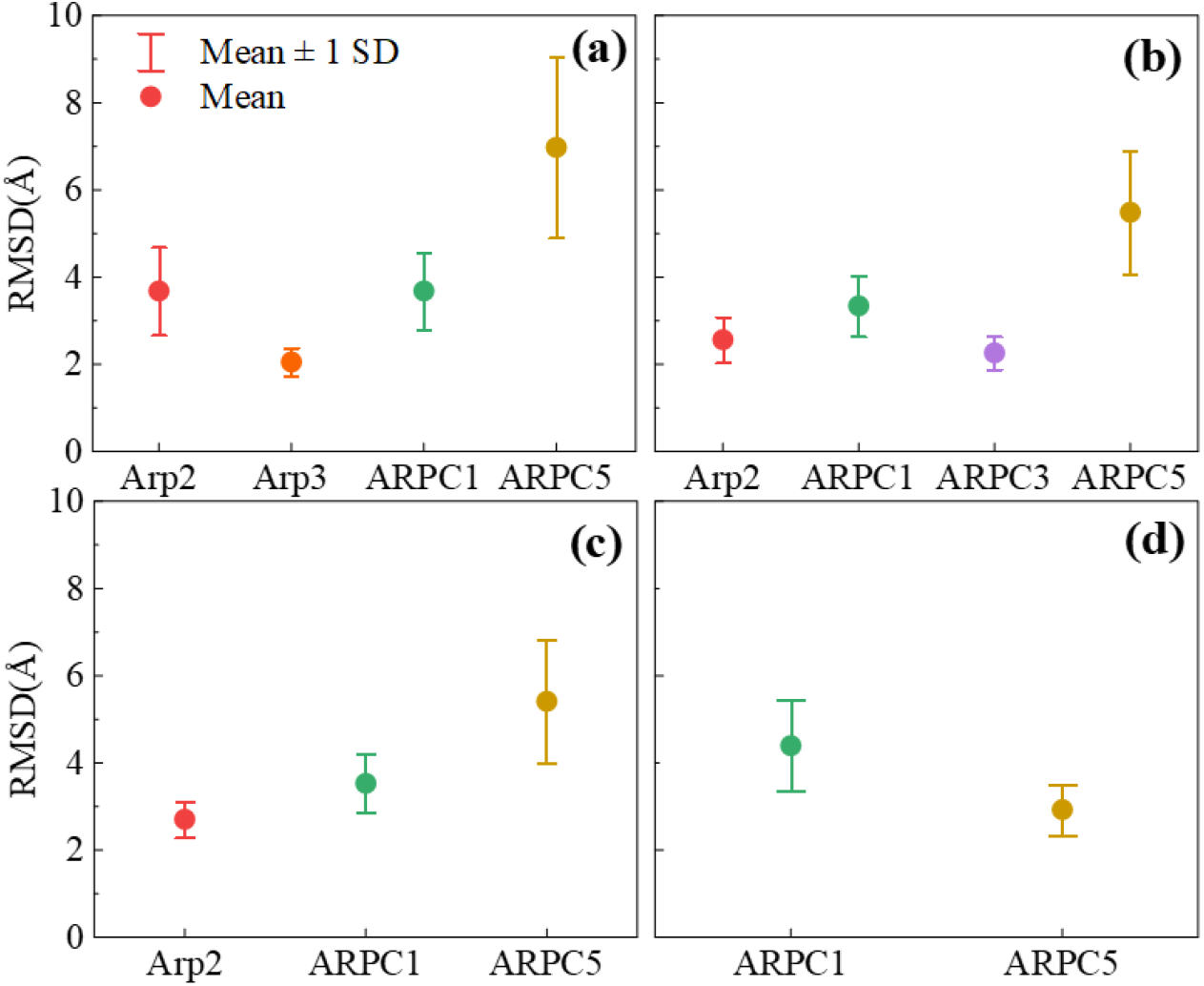
The stability of subcomplexes during the assembly of the Arp2/3 complex. The mean and standard deviation of the RMSD value of each subcomplex after adding a subunit from (a) the second to (d) the fifth iteration. The RMSD value of each CG simulation was calculated using the last 100 conformations of the trajectory.

In the third iteration, Arp2, ARPC1, ARPC3, and ARPC5 were added to the ARPC2-ARPC4-Arp3 trimer, and a CG simulation was subsequently conducted for each tetramer. As shown in Figure 8b, the average RMSD of the subcomplex ARPC2-ARPC4-Arp3-ARPC3 is the smallest, which indicates that this tetramer has the highest stability. ARPC3 may assemble after Arp3.

We repeated the above procedure until a complete protein complex was formed. In the fourth iteration, the most stable five-subunit subcomplex is identified as ARPC2-ARPC4-Arp3-ARPC3-Arp2 (**Fig. 8c**). Arp2 may assemble after ARPC3. Finally, in the fifth iteration, ARPC5 may assemble before ARPC1 because the former can form a relatively more stable six-subunit subcomplex than the latter (**Fig. 8d**). ARPC1 may be the last assembled subunit. Notably, although ARPC1 always has smaller average RMSD values than ARPC5 in iterations 2-4 (**Fig. 8a-8c**), the latter has a smaller RMSD than the former after the addition of Arp2 to the subcomplex (**Fig. 8d**). This may be because ARPC5 can form more native contacts with Arp2 (Table 2), thus increasing the stability of the subcomplex.

Therefore, the predicted assembly order of the Arp2/3 complex is ARPC2, ARPC4, Arp3, ARPC3, Arp2, ARPC5 and ARPC1. Again, this is the inverse of our predicted disassembly order by CG simulations at different temperatures.

The predicted assembly order of the Arp2/3 complex is in agreement with data obtained from some in vitro experimental studies. Reconstitution experiments indicated that ARPC2 and ARPC4 form a structural core of the complex^[39]^, which defines the center of rotation and translation of two subcomplexes. One subcomplex consists of ARPC2, Arp3, and ARPC3, and the other consists of ARPC4, Arp2, ARPC1, and ARPC5.

### Comparison with the method based on interface size

In this section, we compare our results with those obtained from a simple method based on interface size^[18, 19]^. Starting from an intact protein complex, its subunits are dissociated iteratively, and at each step, the subunit with the smallest interface size is removed. The interface size between two subunits is defined by the number of contacted residues, and the map of interface size of a protein complex can be found at the 3D complex server^[20]^. Based on the interface size map of the phosducin-Gtβγ complex, it is possible to predict the disassembly order, which is the phosducin and the Gtβγ dimer. The result is consistent with the disassembly order predicted by our method (**Fig. 4**).

According to the map of interface size of the Arp2/3 complex, a disassembly order from the core subunit ARPC2-ARPC4 is predicted, that is ARPC3, ARPC1, ARPC5, Arp2, and Arp3^[20]^. Compared with the disassembly order predicted by our method, the only difference is ARPC3. The interface size of ARPC3 is the smallest among the seven subunits in the Arp2/3 complex, so it is predicted as the first dissociated subunit. However, in our CG simulations, the disassembly temperature of ARPC3 is 350 K that is significantly higher than those of ARPC1 (315 K), ARPC5 (315 K), and Arp2 (316 K) (**Fig. 6 and 7**).

## Conclusion

Protein complexes play crucial roles in nearly all biological processes. The objective of this work is to examine the order in which subunits assemble and disassemble within a protein complex. We propose a protocol for predicting disassembly processes through CG simulations at multiple temperatures. This technique allows for a comprehensive examination of the disassembly order, comprising definable intermediate subcomplexes. Another protocol is developed that enables prediction of the assembly order. The results obtained for the phosducin-Gtβγ complex and the Arp2/3 complex are in fairly agreement with some previous experimental data.

Despite the simplicity of the CG model, it can still provide helpful insights into the assembly/disassembly of protein complexes. It should be noted that the disassembly temperatures of subunits obtained by CG simulations do not correspond to the actual temperatures, which are used only to judge the disassembly order of the subunits. For a homomeric protein complex, it has been found that the subunits tend to disassemble simultaneously in CG simulations; thus, our method may be more suitable for predicting the assembly/disassembly order of heteromeric protein complexes.

In this preliminary work, only two protein complexes were used to test the protocols. There are certainly some issues to be addressed. We would gather information of more protein complexes, and predict assembly/disassembly orders using our methods and the method based on the interface size. Are the assembly and the disassembly orders of protein complexes are always reversible? Do multiple assembly/disassembly pathways exist? How many different assembly/disassembly orders could be predicted between these methods? Can these differences provide new insight in the assembly/disassembly of protein complexes? In the future, we also hope to expand the applicability of this method to additional biomolecular complexes, such as protein-DNA and protein-RNA complexes, by taking advantage of proper DNA/RNA CG models.

## Acknowledgments

The Supercomputing Center of the USTC provided computer resources for this project, and we are grateful to Mr. Yundong Zhang for his technical support.

